# A cerebellar mechanism for learning prior distributions of time intervals

**DOI:** 10.1101/155226

**Authors:** Devika Narain, Mehrdad Jazayeri

**Affiliations:** Department of Brain & Cognitive Sciences, McGovern Institute for Brain Research, Massachusetts Institute of Technology, Cambridge, Massachusetts 02139, USA

## Abstract

Knowledge about the statistical regularities of the world is essential for cognitive and sensorimotor function. In the domain of timing, prior statistics are crucial for optimal prediction, adaptation and planning. Where and how the nervous system encodes temporal statistics, however, is not known. Deriving from physiological and anatomical evidence for cerebellar learning, we develop a computational model that demonstrates how the cerebellum could learn prior distributions of time intervals and support Bayesian temporal estimation. The model shows that salient features observed in human Bayesian time interval estimates can be readily captured by learning in the cerebellar cortex and circuit level computations in the cerebellar deep nuclei.

---

Human timing behavior is associated with two robust properties. First, response times are biased towards the mean of previously encountered intervals^1^ and second, the bias is stronger for longer sample intervals^2-4^. Bayesian models predict both properties accurately ^2-4^. These models attribute the bias to learning of the prior distribution through previous encounters and predict larger biases for longer intervals, which are more uncertain due to scalar variability in timing ^5,6^. Despite the remarkable success of Bayesian models in describing human timing behavior, little is known about the brain structures and mechanisms that support learning of prior distributions to enable Bayesian integration.

Several considerations suggest that the cerebellum might play a role in learning sub-second to second temporal associations in sensorimotor behavior ^7-12^. First, the rapid learning in behavioral experiments ^13-15^ is consistent with the relatively fast learning dynamics in the cerebellum ^16^. Second, lesions of the cerebellum impact temporal coordination without necessarily influencing movement ability ^17^. Third, human neuroimaging experiments implicate the cerebellum in timing ^18,19^. Fourth, work in non-human primates suggests that the cerebellum is involved in a range of sensorimotor and non-motor timing tasks, from temporal anticipation during smooth pursuit^20,21^ to detecting oddballs in rhythmic stimuli^22^, to timing self-initiated movements ^23^. Finally, studies of eyeblink conditioning in humans as well as numerous animal models ^24^ suggest that the cerebellum is one of the key nodes involved in learning the interval between conditioned and unconditioned stimuli.

Cerebellar circuits can learn multiple time intervals simultaneously. For example, in eyeblink conditioning, animals can concurrently acquire differently timed conditioned eyelid responses associated with distinctive conditioned stimuli ^25^. The ability to acquire more than one interval suggests that the cerebellum might have the capacity to learn a range of previously encountered intervals. This is an intriguing possibility as it suggests that the cerebellum might play a functional role in Bayesian computations that rely on knowledge about the prior probability distribution of time intervals. Here, we propose a theoretical model called TRACE (Temporally Reinforced Acquisition of Cerebellar Engram) that synthesizes known anatomical and physiological mechanisms of the cerebellum to explore how prior distributions could be encoded to produce Bayesian estimates of time intervals.

## Results

### Behavioral paradigm

To assess the potential role of the cerebellum in Bayesian time estimation, we focused on a simple time interval reproduction task (Fig. 1a). In this task, which we will refer to as the *Ready-Set-Go* (RSG) task ^2,26^, two cues, *Ready* and *Set*, demarcate a sample interval drawn from a prior distribution, that subjects estimate and subsequently reproduce. Previous work has shown that both humans and monkeys exhibit the two classic features of Bayesian timing while performing this task (Fig. 1b): produced intervals are biased towards the mean of the prior distribution, and this bias is larger for longer and more uncertain intervals ^27,28^. We use this task to examine whether and how the cerebellum could acquire the prior distribution of time intervals and compute a Bayesian estimate of the measured interval.

**Figure 1:**
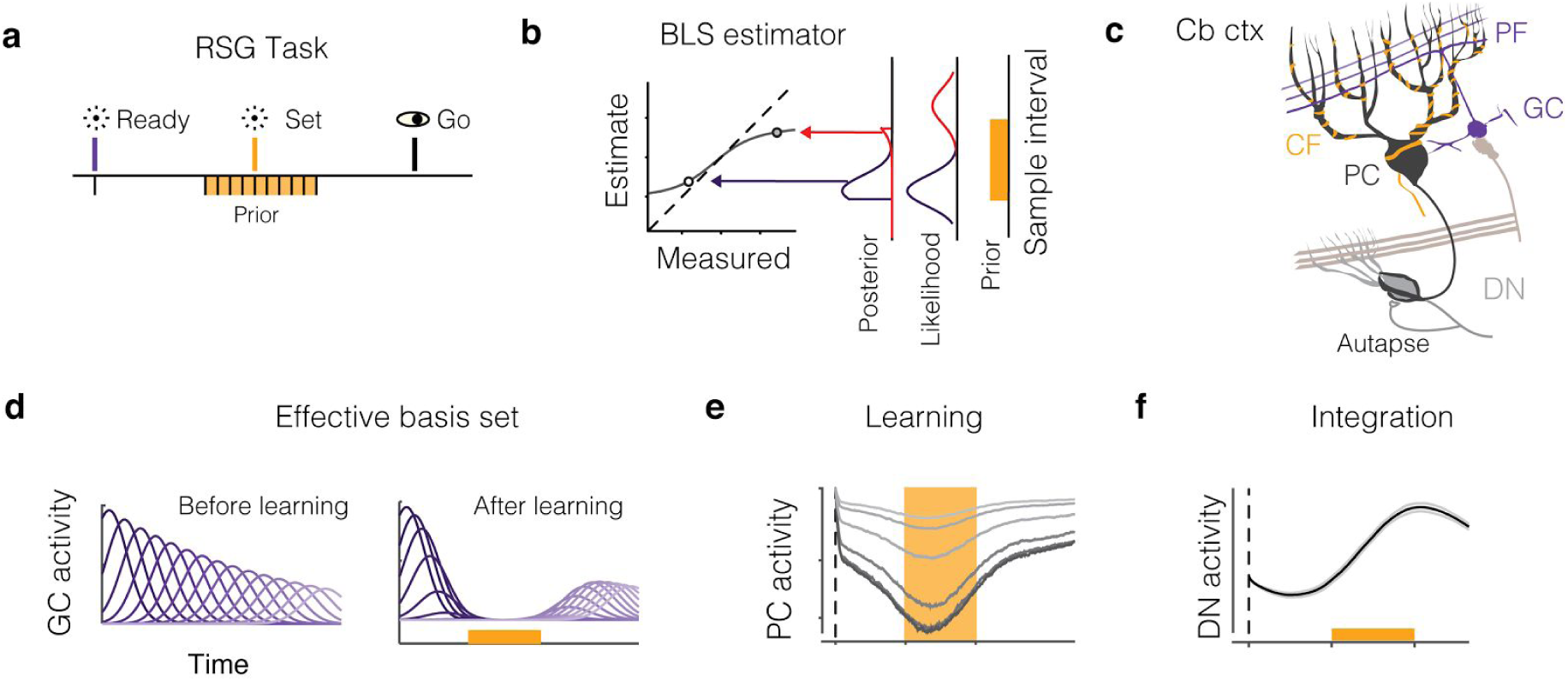
Task, anatomy and the TRACE model. a) Ready-Set-Go task. On each trial, subjects measure a sample interval demarcated by two visual flashes – Ready followed by Set – and reproduce it immediately after Set. The sample interval is drawn from a uniform prior distribution (orange), b) Bayes-Least-Squares (BLS) estimator. A Bayesian observer computes the posterior based on the product of the prior and the likelihood function, and uses the mean of the posterior to estimate the interval. The plot shows the behavior of BLS for two trials with a uniform prior. The product of a uniform prior with the likelihood function associated with the two measurements (red and blue bell-shaped curves) truncates the likelihood functions such that the resulting mean is biased toward the mean of the prior distribution (arrows). This BLS is characterized by a nonlinear function that biases the estimates towards the mean of the prior. For longer measurements (e.g., red compared to blue), due to scalar variability, the likelihood is wider, which causes the magnitude of the bias to be larger, c) Schematic drawing of the cerebellum. In the cerebellar cortex (Cb ctx), a Purkinje cell (PC, black) receives inputs from granule cells (GC, purple) via parallel fibers (PF), and climbing fibers (CF, orange). PCs, in turn, project to and inhibit neurons in the dentate nucleus (DN), which additionally receives extra-cerebellar input (light brown) and autaptic (synapse onto self) input, d-e) Components of the TRACE model, d) The effect of the basis set on PCs (effective basis set). Left panel shows the basis set activity (sub-sampled from N = 500) before learning, and the right, shows GC activity filtered by the GC to PC synapses after learning, e) Changes of PC activity profile during learning. The PC activity is shown after 10, 50, 100, 150 and 200 trials during learning (light to dark), f) DN activity. The activity profile ofDN reflects the integrated output after learning (trial 200). Error bars indicate SEM.

### TRACE model

The TRACE model consists of three components, which are motivated by the known anatomy and physiology the cerebellum (Fig. 1c). The first of these is a *basis set*, that represents reproducible and heterogenous patterns of activity across granule cells (GCs, Fig. 1d). Second is *learning*, which relies on known plasticity mechanisms at the synaptic junction between GC axons (parallel fibers PF) and Purkinje cells (PCs, Fig. 1d, right and 1e). Third is integration, which captures transformations downstream of the cerebellar cortex (Fig. 1f).

#### Basis set

The first component of TRACE is a heterogeneous temporal basis set across GC neurons (Fig. 1d). Recent studies have provided convincing evidence for the existence of such heterogeneity ^29,30^, and models of cerebellar tasks have highlighted the computational advantage that such a basis set could confer upon the learning of temporal contingencies ^31–34^.

Similar to previous work ^32,33^, we assumed that the *Ready* cue would trigger the basis set across GCs. We modeled this basis set as Gaussian kernels across time (Supp. Fig. 1a). A large body of literature in animal and human studies suggests that the representation of time in the brain is subject to scalar noise; i.e., noise whose standard deviation scales with elapsed time ^6^_’_^35^. We used a probabilistic model to characterize the effect of scalar noise on the expected profile of the basis set (Supp. Fig. 1b). The model demonstrated that scalar noise distorts the expected profile of the kernels in two ways: it would cause amplitudes to decay and widths to increase with time. These effects were accurately captured by augmenting the model of the basis set rates (*r*(*t*)) with an exponential decay *g*(*t*) in amplitude and a linear increase in width (*σ_basis_*) with time (Fig. 1d, Supp. Fig. 1b; see Methods).

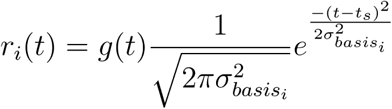

where

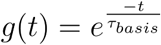

#### Learning

The acquisition of time intervals in the cerebellar cortex is primarily attributed to synaptic changes between GCs and PC neurons. These synapses are thought to undergo long-term depression and potentiation (LTD and LTP) ^36–39^. Depression of synaptic weights through LTD depends on the conjunctive activation of GCs and complex spikes triggered by climbing fibers (CFs), whereas LTP occurs when GCs are active in the absence climbing fiber activity. Learning in TRACE therefore depends on these two complementary mechanisms.

We assumed that *Set* activates CFs and causes LTD in the subset of synapses that are activated by GCs shortly before the time of *Set.* Since the activity of GCs is triggered by *Ready*, this plasticity mechanism causes an interval-dependent decrease in PC activity that reflects the previous *Ready-Set* interval. The LTP, on the other hand, potentiates those GC to PC synapses associated with the subset of GCs that are active in the absence CF activity. The learning rule we used is similar to previous modeling work on LTP and LTD learning in the cerebellum^40^ with the following specifications:

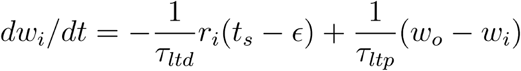

LTD (first term) was activity-dependent (dependent on *r_i_*(*t*)), and effective only for eligible synapses. A synapse was considered eligible if the corresponding GC was active within an eligibility trace (*ϵ*) and based on previous reports for eyeblink conditioning ^32,41^, we estimate to be 50 ms. In contrast, LTP (second term) acted as a restoring force driving the synaptic weight towards a baseline (*^ɷ^o*) when GCs fired in the absence of CF stimulation. This learning rule was governed by two free parameters: the time constants of LTP and LTD (*τ_ltd_* and *τ_ltp_*).

The LTD component of this learning rule permits each presented sample interval to reduce the synaptic weight of the subset of GCs in the basis set that are *eligible* at the time of Consequently, multiple exposures to sample intervals drawn from a prior distribution (Fig 1a, orange), allow GC-PC synapses to gradually acquire a representation of the full prior distribution (Fig. 1d, right). LTP, on the other hand, gradually washes out LTD allowing learning to become adaptive. More specifically, LTP allows synaptic weights to have a stronger footprint for intervals that are presented more frequently. LTP also allows synapses to represent the most recently encountered time intervals. The speed with which the model adapts to changes depends on the relative time constants associated with LTP and LTD. We demonstrate that the final output of TRACE is relatively robust to variations of these time constants (Supp. Fig 4).

Modification of GC to PC synapses directly impacts PC activity, since it represents the net granule cell activity weighted by the synaptic weight. We modeled PC activity (*V_pc_*) as the linear sum of GC basis set activity filtered by the GC-PC synaptic weights. Accordingly, PC activity is influenced by both the response profile of the GC basis set and the learned synaptic weights (Fig. 1e left). This enables PCs to encode a composite variable that carries information about both uncertainty in the measurement (via the basis set) and prior distribution (via the synaptic weights).

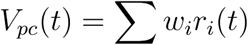

An implicit assumption of this learning scheme is that the system must correctly route the stimuli such that *Ready* would activate GCs and *Set* would activate CFs. However, such precise routing of information is not necessary, and TRACE can straightforwardly accommodate a situation in which *Ready* and/or are also present on the alternate pathway. This flexibility can be understood by considering the effect of the eligibility trace ^42,43^ on cerebellar LTD. The eligibility trace would render activation of CF by *Ready*, at the start of the interval, inconsequential as there would be no systematic GC basis set activity prior to *Ready.* The activation of GCs by *Set*, after the interval would be similarly inconsequential as there would be no CF activity to cause LTD. Therefore, the primary LTD-dependent learning in TRACE would occur when Set-dependent CF activation causes LTD in eligible GC to PC synapses.

#### Integration

The last stage of TRACE is concerned with the transformation of PC activity in the cerebellar deep nuclei (Fig. 1f). We focused our attention on the caudal region of the dentate nucleus where timing signals in both motor ^23^ and non-motor timing tasks ^22^ have been observed at the level of individual dentate neurons (DNs). This region contains large columnar neurons with three major synaptic inputs: extra-cerebellar currents, inhibition from PCs, and autaptic currents ^44^. Accordingly, the membrane potential of DNs can be modeled as follows:

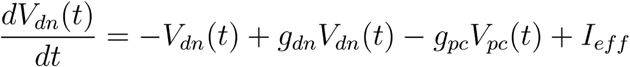

where *g_dn_* and *I_eff_* correspond to the conductance associated with the autaptic input, the conductance associated with PCs, and the remaining effective input driving DNs.

In this model, the transformation of *V_pc_*(*t*) by the DNs depends on two main factors, the autaptic conductance (*g_dn_*), and the effective input drive (*I_eff_*). As has been shown by previous work on autaptic architectures, *g_dn_* currents counteract the leakage current and allow the neuron to act as an integrator^45^. For the remainder of the manuscript, we assume that *I_eff_*, which is a constant positive drive, is equal to the average of *g_pc_V_pc_*(*t*) over time. We later show that relaxation of this assumption does not impact the model behavior (Supp. Fig. 2a-c). We also assume that DNs act as perfect integrators (i.e., *g_dn_* = 1) and integrate the inputs from PCs and the input current ( *I_eff_* – *g_pc_V_pc_*(*t*)). We also show that the model is robust to changes in the value of *g_dn_* (Supp. Fig 2d-f). With these assumptions, the membrane potential of the DN can be computed as follows:

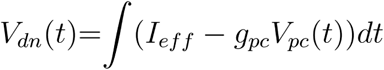

**Figure 2:**
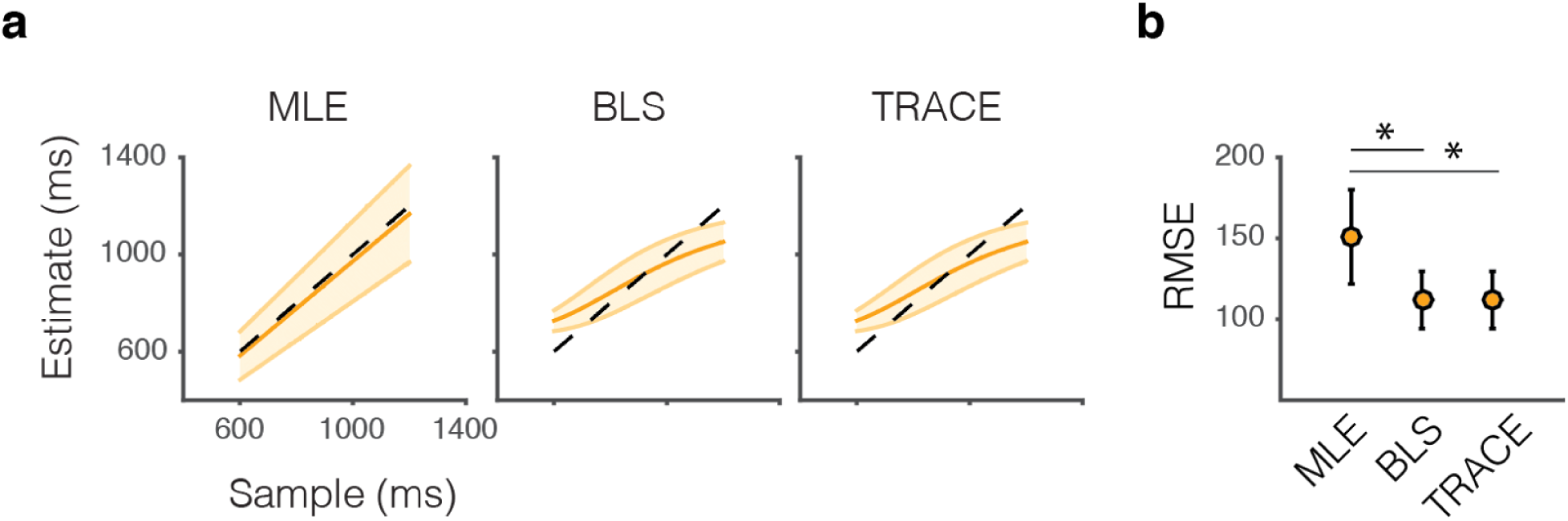
Bayesian estimation by TRACE. a) The maximum likelihood estimator (MLE). The plot shows the behavior of the MLE in the RSG task. MLE is suboptimal as it does not take the prior distribution into account, b) The Bayes least-squares (BLS) estimator. The BLS provides an optimal estimate as it takes full advantage of both the likelihood function and the prior distribution (Fig.1b), c) The output of TRACE. TRACE quantitatively matches the behavior of BLS. Error bars (shaded area) represent SEM. b) Root-mean-squared error (RMSE) between estimates and sample intervals for MLE, BLS and TRACE. Asterisks indicate p << 0.001. Error bars indicated standard deviation.

### Bayesian estimation by TRACE

We compared the output of TRACE with the Bayes-Least-Squares (BLS) estimator, which accurately describes human behavior in the RSG task ^2^. BLS acts as a nonlinear transformation of a measured variable to an optimal estimate (Fig. 1b) and has two key characteristics: 1) estimates are biased towards the mean of the prior, and 2) due to scalar variability, the magnitude of the bias is larger for longer intervals. As shown in Fig. 2, the integrated output of the TRACE (i.e., the DN activity) accurately captured the nonlinearities associated with BLS.

We compared the RMSE for BLS, TRACE and a sub-optimal maximum likelihood estimator (MLE) (Fig. 2). Both the BLS and TRACE outperformed the MLE (paired t-test BLS-MLE: t_399_ = 30.312, p << 0.001 TRACE-MLE, t_399_ = 30.205, p << 0.001) and there was no significant different between the BLS and the TRACE (paired t-test t_399_ = 0.047, p = 0.961). Therefore, TRACE captures both temporal uncertainty and prior-dependent biases in a manner consistent with Bayesian estimation theory. Finally, the Bayesian behavior of TRACE is robust to variation of parameters in the basis set and learning rate parameters (Supp. Fig. 3-4).

**Figure 3.**
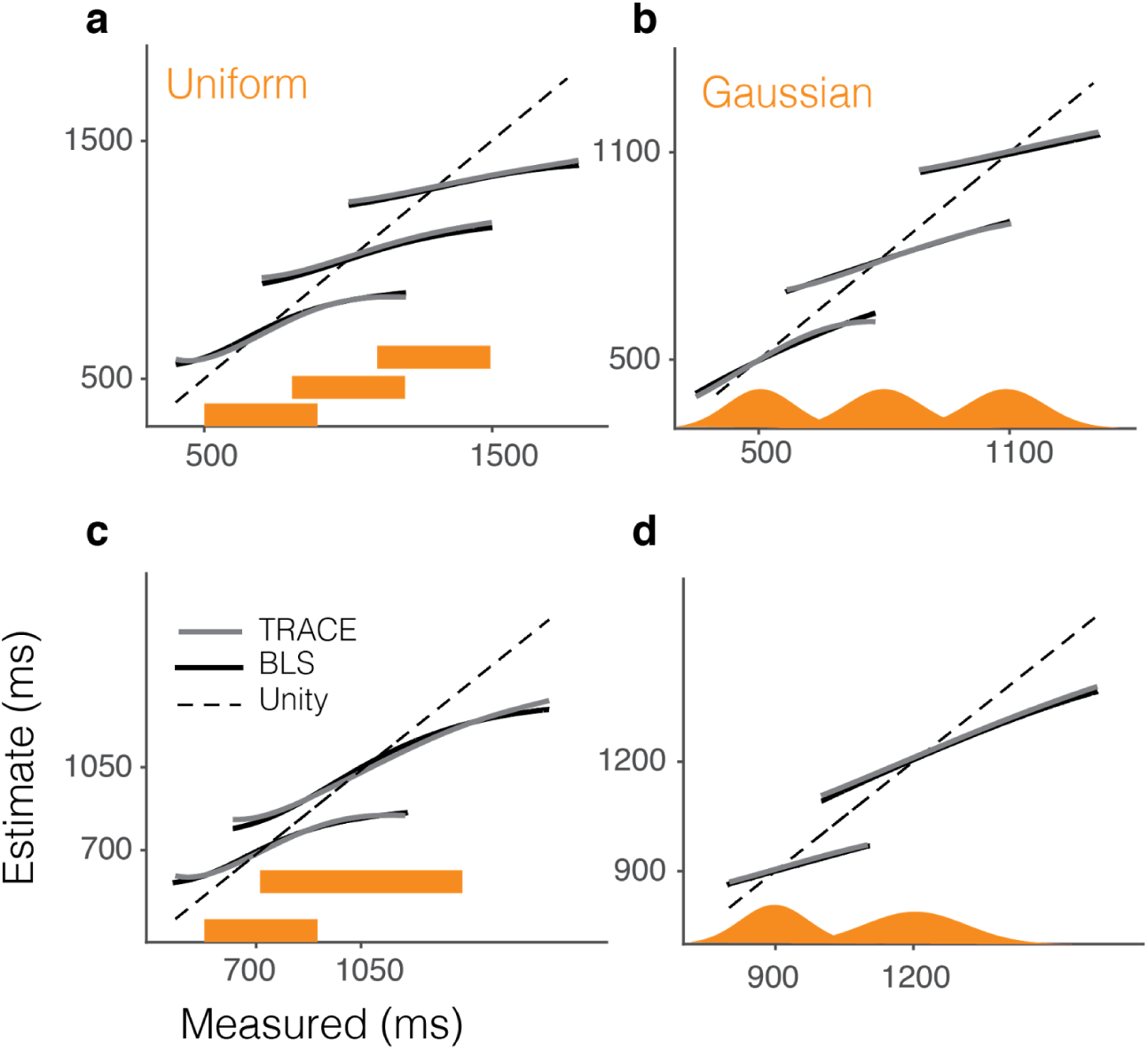
Comparison of TRACE and BLS for different priors. a) Estimates derived from Bayes least-squares (BLS) (black line) and TRACE (gray line) for three different uniform prior distributions (orange) with different means, b) Same as a) for three Gaussian prior distributions, c) Same as a) for two uniform priors with different means and widths, d) Same as c) for Gaussian priors. We used a linear scaling parameter to calibrate the output of TRACE and added the mean of the prior as an offset so that TRACE and BLS could be compared directly.

**Figure 4.**
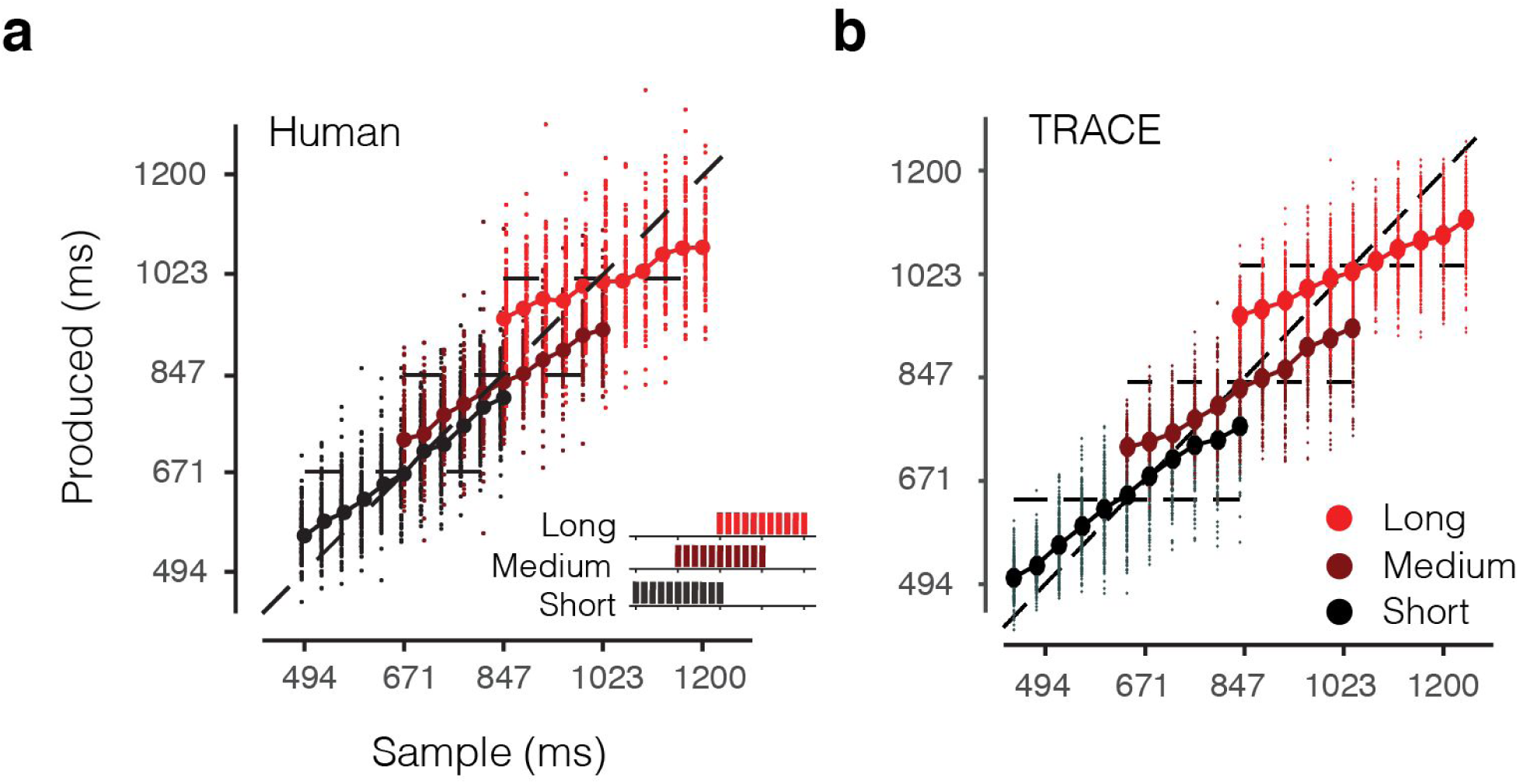
Comparison of TRACE with human behavior in the RSG task. a) Human behavior during the RSG task with three different prior distributions, short (black), medium (crimson) and long (red). Data adapted ^2^. b) The corresponding prediction from TRACE. Dots and circles mark individual responses and averages respectively. Parameters of the model were not tuned to the data and a single linear scaling factor was used to calibrate model output for all conditions ^2^.

Next, we tested whether TRACE could perform Bayesian estimation in the presence of uniform and Gaussian distributions with different means and standard deviations (Fig. 3). Remarkably, the output of TRACE matched the behavior of a BLS estimator across all conditions without any need to adjust model parameters. In the case of uniform priors with different means, we compared the behavior of TRACE to the behavior of human subjects reported previously ^2^ and found that the model was able to accurately capture the observed biases (Fig. 4).

### TRACE learning dynamics

The learning dynamics in TRACE can be evaluated in terms of two variables, the asymptotic change in synaptic weights and the effective time constant at which synapses reach that asymptotic value. Both the asymptotic weight change and the effective time constant are influenced by the time constants associated with LTP to LTD (Supp. Fig. 4). Overall the *τ_ltp_* has to be larger than *τ_ltd_* for stable learning to occur. As *τ_ltp_* becomes increasingly larger, the model establishes a stronger and more resilient footprint of previously encountered time intervals (larger weight change). This however, comes at the cost slower forgetting of learning and as a result, slower adaptation to recent changes. The effective time constant also varies with both *τ_ltd_* and *τ_ltp_* (Supp. Fig. 4).

We tested the model’s ability to capture the dynamics of learning prior distributions in a previous behavioral study involving a time reproduction task similar to RSG ^13^. In that study, subjects gradually adjusted their behavior when, unbeknownst to them, the prior distribution was altered. Interestingly, the adjustments were slower when the prior switched from wide to narrow compared to vice versa (Fig. 5a). Similar results for switching between wide and narrow priors have been reported in other Bayesian tasks in the sensorimotor domain ^46,47^. TRACE exhibited the same behavior as evidenced in the changes of weights in the GC-PC synapses: learning was relatively slower after switching from a wide to a narrow prior (Fig. 5b). This behavior can be understood by comparing the action of LTP and LTD after the two kinds of switches. When the prior switches from wide to narrow, LTP restores the depression associated with intervals that are no longer presented. In contrast, when the prior switches from narrow to wide, LTD creates a footprint for the newly presented intervals. Since LTP is slower than LTD (to help retain information from past trials more effectively), learning in the former conditions proceeds more slowly.

**Figure 5.**
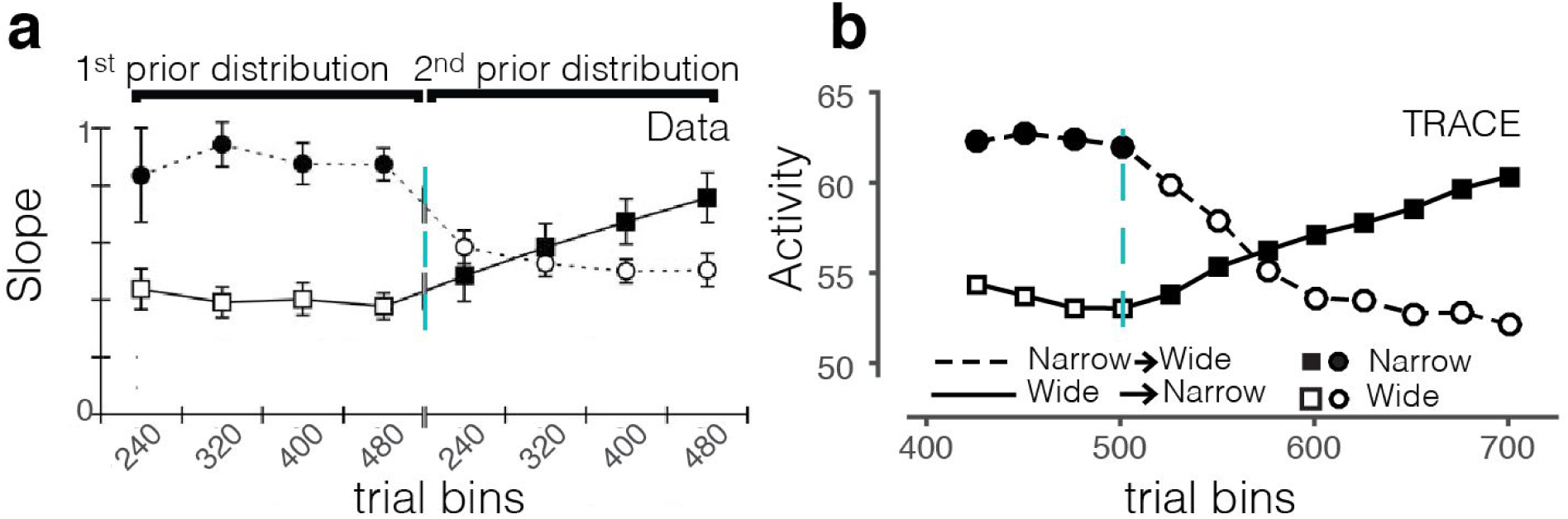
Comparison of TRACE with human behavior during learning. a) Human behavior during transitions between a wide and narrow prior distribution in a time interval reproduction task similar to RSG. Data is adapted ^13^. The results were quantified by a ‘slope’ parameter that quantified the strength of the regression towards the mean of the prior (higher slope represents more bias). The switch (cyan) from narrow (filled symbols) to wide (open symbols) decreases the influence of the prior and vice versa, b) The corresponding prediction from TRACE (see Methods). The change in PC activity compared to baseline is plotted on the ordinate (in arbitrary units). TRACE parameters were the same as those used in RSG and were not fit to the data.

### Comparing TRACE output to neural activity

In TRACE, DN neurons integrate the activity of PCs across time. This makes a specific prediction for how the output of the cerebellum (i.e., DN activity) would reflect previously encountered time intervals (i.e. prior distribution) in RSG. No experiment has recorded from DN neurons in RSG, but a recent study characterized activity in the lateral intraparietal cortex (LIP) of monkeys’ in a variant of the RSG task with a uniform prior ranging between 529 and 1059 ms ^28^. Since LIP neurons receive transthalamic input from DNs ^48,49^, we compared predictions of the model to LIP activity. LIP neurons were modulated throughout the *Ready-Set* epoch and displayed a nonlinear response profile during the support of the prior distribution (between 529 and 1059 ms). This profile was highly similar to the output of TRACE (Fig. 6). This similarity is consistent with TRACE being involved in Bayesian integration of time intervals and suggests that the nonlinear response profiles observed in LIP may be inherited from the output of the cerebellum.

**Figure 6.**
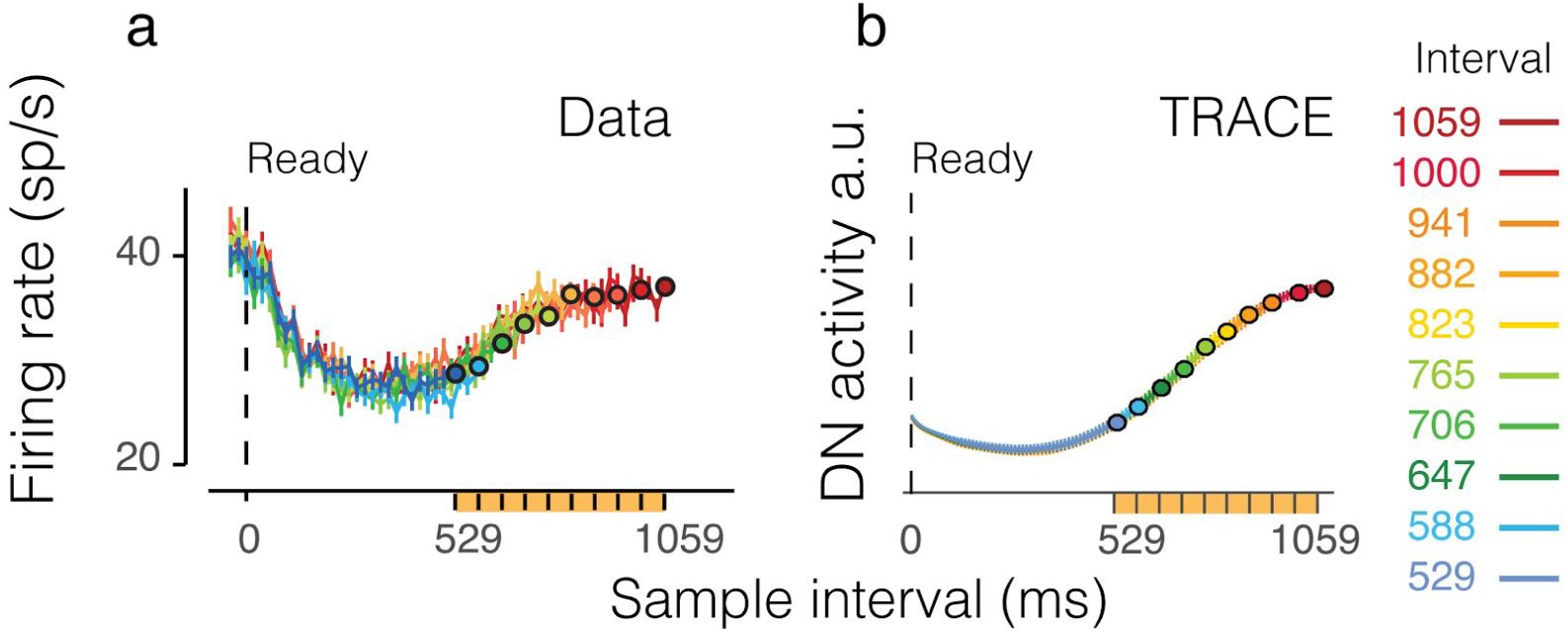
Comparison of TRACE with electrophysiology data. a) Physiology. Activity in the lateral Intraparletal (LIP) area in monkeys performing the RSG task is modulated nonlinearly during the range of the prior (orange) between 529-1059 ms. Data adapted from ^28^. b) The corresponding prediction from TRACE. The output of the TRACE is shown with arbitrary units. Error bars indicate SEM.

### Model lesions

To evaluate the three key components of TRACE (basis set, learning and integration), we examined its behavior after ‘lesioning’ each of these components. For the basis set, we first simulated a variant of TRACE in which the basis set did not decay with time (Fig. 7b). This disruption led to a loss of the interval-dependent asymmetry in the bias (i.e., stronger bias for longer intervals), which is a key feature of Bayesian integration. In contrast, if the basis set decays too rapidly, the model fails to capture prior-dependent biases for longer intervals (Fig. 7c). The learning was also critical. When no learning was allowed, TRACE was insensitive to the prior distribution (Fig. 7d). Finally, without the integration component, the time course of the model output did not reflect the monotonic increase of human Bayesian estimates with interval duration (Figure 7e). These analyses validated that all three components of the model were necessary for the induction of Bayesian behavior.

**Figure 7.**
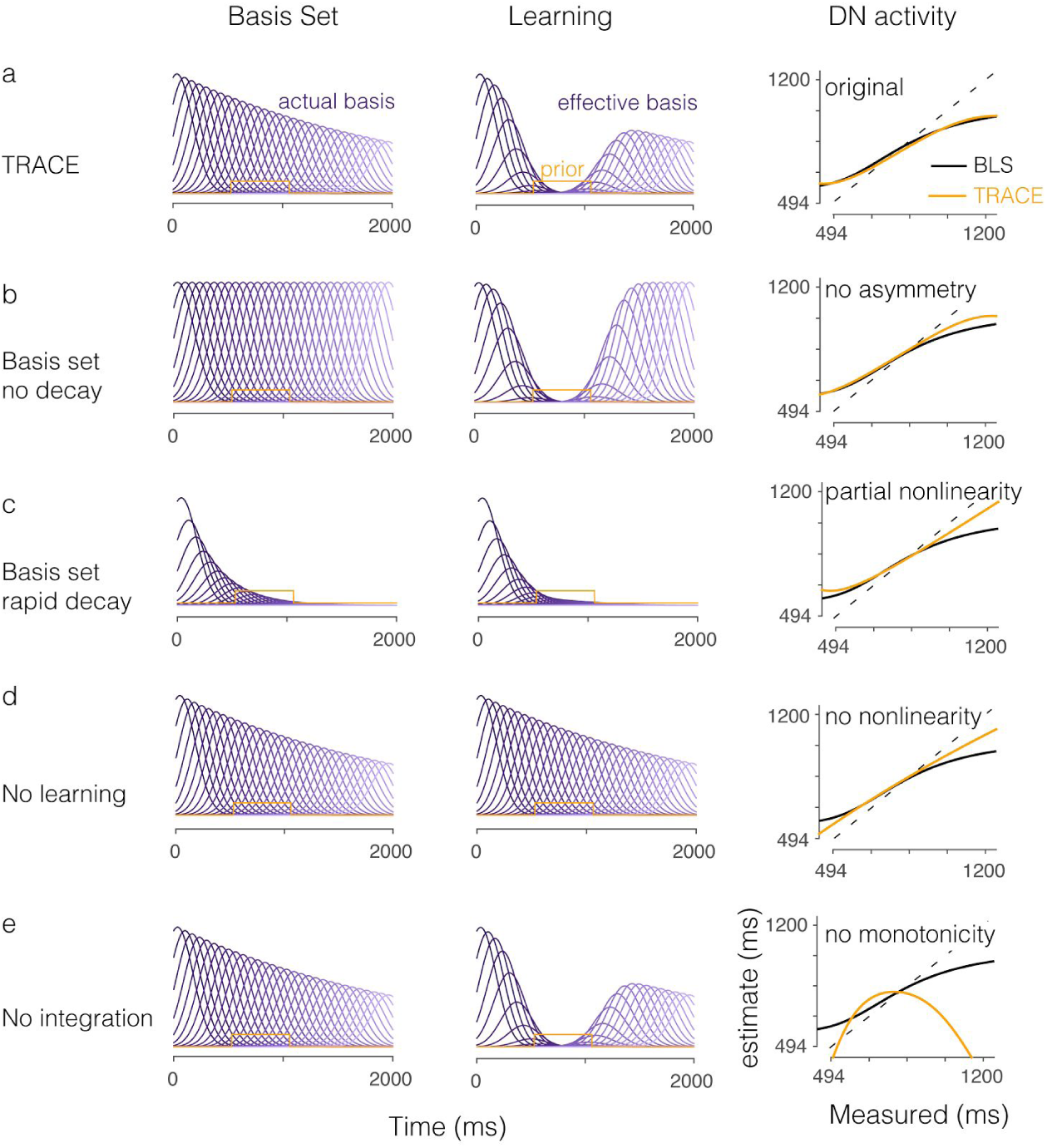
TRACE lesions. a) TRACE with all three model components intact. Left: An attenuating and widening basis set of granule cells. Middle: The effective influence of the basis set after learning of the prior distribution has reached steady state. Right: The linearly transformed output of TRACE (orange) with the BLS function (block). b) Same as for a. The asymmetry in TRACE output is lost. c) Same as earlier panels. Due to lack of temporal elements for longer intervals. no nonlinearity is acquired for these. d) If no learning takes place, then there are no significant biases in the model output towards the mean of the prior, signalling that while the ability to do the task is not compromised in the absence of learning in TRACE, the ability to do so in a statistically optimal manner, as humans do, is los., e) If no integration takes place, there is no monotonically increasing output, leading to significant deteriorations in performance. Therefore the model predicts that lesioning the deep nuclei will cause severe problems with performing the task accurately, whereas lesions to the cortex will lead to more subtle behavioral differences.

## Discussion

Numerous behavioral studies have shown that humans rely on prior knowledge to mitigate the uncertainty in sensory measurements, as predicted by Bayesian integration theory^2–4,13,46,50–52^. This raises the possibility that brain circuits have the capacity to encode prior distributions and use them to optimize behavior. However, where and how prior distributions are represented in the brain is a matter of debate. In sensory domains where prior knowledge is characterized by the natural statistics of the environment, it is though that the priors are encoded implicitly by the organization of synaptic connections in sensory areas ^53–56^. In sensorimotor and cognitive tasks, Bayesian inference is thought to occur later in the association and premotor cortex ^28,57–61^. In the domain of time, there is strong evidence that humans rapidly learn and utilize prior distributions of time intervals to optimize their performance^2–4,28,62,63^. In this study, we asked what brain structure could suitably encode prior distribution of time intervals within the sub-second to seconds range, which is also crucial for sensorimotor behaviors.

Two general lines of reasoning led us to hypothesize that the cerebellum may be a key node for Bayesian timing. First, converging evidence from human patients and imaging to neurophysiology experiments in various animals models suggests that the cerebellum plays a central role in timing tasks ^8,17,22,64,65^ Second, the cerebellum is thought to be involved in acquiring and updating internal models of movement ^66,67^, which implies that the cerebellum has the capacity to learn the temporal contingencies that relate sensory inputs to motor outputs and vice versa. This led to the examination of whether the cerebellum could support learning distributions of time intervals and support Bayesian timing.

We constructed a model based on the known anatomy and physiology of the cerebellum with three specific components, a basis set, a learning rule, and an integration stage. The basis set was inspired by the recent studies indicating that GC have temporally heterogeneous activity patterns ^29,30,68^. We modeled this by assuming that GCs form a temporal basis set composed of Gaussian kernels. This is a simplification as each GC is likely to be activated at multiple time points and the detailed temporal structure of the basis set is likely to depend on task-dependent extra-cerebellar inputs from mossy fibers. Nonetheless, considering the massive convergence of GCs to PCs, coupled with the heterogeneous activity patterns of each GC suggests that we can model the input to each PC by a temporally dense basis set. A key assumption in the construction of the basis set was that GC activity is subject to cumulative temporal noise resulting in trial-by-trial jitter in GC activity. What motivated this assumption was the presence of scalar variability in behavior, which is arguably among the most consistent features of timing across tasks and species ^6,69,70^. We did not attempt to develop a biophysical model capturing this variability as the generative process underlying scalar variability was not relevant to our objective (i.e., learning of prior distribution and Bayesian integration). However, we verified that increased variability with time would emerge naturally within a noisy recurrent network (Supp Fig. 5). This assumption can be verified experimentally by evaluating whether trial-averaged activity of granule cells attenuates over time and becomes more variable.

The learning rule in TRACE consists of LTD that weakens eligible GC to PC synapses upon the activation of CFs, and LTP, which restores synaptic weights in the absence of CF activity. Although this formulation is relatively standard, complementary plasticity mechanisms might also be at play as recent work has begun to demonstrate ^71,72^. For example, plasticity mechanisms in GC to PC synapses may be tuned to diverse intervals and may be region and task dependent ^73^ providing complementary substrates for learning time intervals. Similarly, certain aspects of learning are thought to depend on intracellular PC mechanisms ^74,75^. Therefore, future work might seek to augment the standard learning rules in TRACE based on the relevance of alternative plasticity mechanisms for learning time interval distributions.

In the context of RSG, we examined the consequence of *Ready* and *Set* visual flashes activating GCs and CFs, respectively. We did not consider the effect of *Ready* and *Set* on the alternative pathways, which as we described previously, is inconsequential to the behavior of TRACE. There are numerous lines of evidence suggesting that both GCs and CFs receive visual signals. For example, anatomical studies have shown that the LGN ^76,77^, the superficial, intermediate, and deep layers of the superior colliculus (SC)^77,78^, and the visual cortex ^77,79^ project to different regions of the pontine nuclei. Moreover, stimulation of the LGN, superior colliculus or visual cortex is sufficient as a conditioned stimulus to induce eyeblink conditioning ^80^. Therefore, anatomically, there are numerous pathways for visual flashes to drive GCs. Visual input may also be relayed to the cerebellar cortex by the inferior olive through visual afferents such as the superior colliculus ^81,82^. Consistent with this view, visual flashes have been shown to evoke complex spikes in the lateral cerebellum ^83^.

The last component of the model is the integration of PC activity in DN. DN, which is perhaps the least studied output structure of the cerebellum may be particularly important in conveying timing information in the context of non-motor tasks. For example, recent physiology work suggests an intriguing role for DN in a temporal oddball detection task ^22^, which goes beyond traditional motor functions attributed to the cerebellum. A potential non-motor function for DN was also noted by anatomical studies showing that DN interacts with higher cortical areas that do not directly drive movements ^48,84_’_85^. of particular relevance to our work is the interaction between DN and the parietal and supplementary motor cortical areas ^86-90^. These areas have been implicated in a range of timing tasks ^28,91–95^, and the activity of parietal cortical neurons in the context of the RSG task matches the predictions of TRACE remarkably well (Fig. 6).

Our assumption that DN neurons integrate PC activity was motivated by the autaptic organization of large columnar neurons in DN^44^. As numerous theoretical studies have shown, this organization is uniquely well-equipped to carry out temporal integration ^45,96_’_97^. Indeed, the match between the output of TRACE and LIP activity in the RSG task (Fig. 6) as well the ability of TRACE to emulate Bayesian integration (Fig. 2 and 4) was afforded by the assumption of integration. Our model therefore, makes a general prediction that certain DN neurons integrate the signals they receive from both PCs and other extra-cerebellar inputs. Stated differently, TRACE predicts that PC activity carries a time derivative of the cerebellar output provided by DN. This idea is consistent with observations that PCs carry other signals that may represent the time derivative of behavioral outputs such as the speed of hand movements ^98,99^, speed of saccades ^100^, speed of smooth pursuit ^101^, and speed of eyelid in eyeblink conditioning ^102^. We note however, that this integration may also occur further downstream in parietal or frontal cortical areas ^103^_’_^104^, which receive transthalamic input from DN^48,49,105^ and have been implicated in temporal integration of information^28_’_103_’_106^.

TRACE provides a circuit-level description of how the brain could acquire and utilize prior distributions of time intervals. The underlying computation however, does not rely on explicit representations of the traditional components of Bayesian models including the prior, the likelihood and the posterior. Instead, the output of the model represents an online estimate of elapsed time that derives from a composite representation of the prior and the likelihood within the cerebellar cortex. This is akin to establishing a nonlinear transformation that directly maps measured elapsed time to its Bayesian estimate, as was hypothesized previously^2^.

Although we found the circuitry within the cerebellum as particularly suitable for learning prior distributions of sub-second to second time intervals, it is almost certain that the full richness of temporal information processing depends on coordinated interactions of the cerebellum with other brain areas including cortex, the basal ganglia and hippocampus. For example, in the RSG task, evaluation of behavioral performance is likely to engage higher cortical areas ^107^, and the processing of feedback on timing performance is likely to engage the basal ganglia ^108^. Furthermore, numerous studies have found neural correlates of interval timing across the cortico-basal ganglia circuits ^93,109–113^. indeed, even within the context of the RSG task, cortico-basal ganglia circuits are likely to be involved in the conversion of a Bayesian estimate computed at the time of *Set* to an ensuing motor plan for the reproduction of that interval ^114^. Finally, learning of time intervals in the cerebellum may depend on interactions with and inputs from other brain areas. For example, although lesioning of the deep nuclei may cause impairment in both delay and trace eyeblink conditioning ^115^_’_^116^, trace conditioning depends on other brain structures including the hippocampus that is thought to help establish a memory of recently-acquired trace intervals ^117,118^. One possible role of extra-cerebellar inputs might be to help maintain a certain level of activity to facilitate the acquisition of the trace period ^119^. A similar input may be needed for activating the basis set in the RSG task.

In sum, our work highlights the potential for an exciting new function for the cerebellum: the ability to represent and integrate prior distributions of time intervals. Remarkably, this function emerges naturally from what is known about the anatomy and physiology of the cerebellum in the context of tasks that expose the cerebellum to a range of time intervals including the RSG task. The simplicity and success of TRACE in capturing Bayesian theory (Fig. 2), Bayesian behavior (Fig. 4), Bayesian learning (Fig. 5) and related neural responses (Fig. 6) invites a serious consideration of the possibility that the cerebellum is a key component of the circuits that the brain uses to emulate probabilistic computations in the domain of time.

## Methods

### Bayesian estimator

The *Ready-Set-Go* (RSG) time interval estimation task consists of two consecutive cues, and *Set* that mark an interval (*t_s_*) drawn from a prior distribution *π*(*t_s_*). Subjects measure *t_s_* and subsequently reproduce it (Figure 1a). Following previous work^2^, we modeled the measured interval, *t_m_* as drawn from a Gaussian distribution centered at *t_s_* whose standard deviation is proportional to *t_s_* with coefficient of variation *ɷ_m_*.

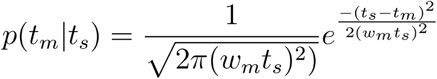

The Bayes-Least-Square (BLS) estimate *t_e_* is the expected value of the posterior distribution *p*(*t_s_*|*t_m_*), as follows:

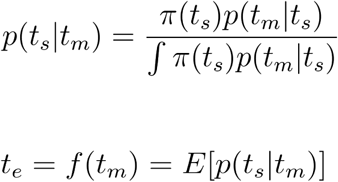

where *E*[.] denotes expectation. The BLS estimator can be formulated as a deterministic nonlinear function, *f*(*t_m_*), that maps a noisy *t_m_* to an optimal estimate *t_e_* (Figure 1b). In the manuscript, we assessed the behavior of the BLS estimator for different uniform and Gaussian priors and for *ɷ_m_* = 0.1. This value is consistent with previous reports for humans performing the RSG task^2^.

### TRACE model

In TRACE, Purkinje cells (PCs) receive input from *N* granule cells (GCs). The spike count (*y*) for the *i^th^* GC follows an inhomogeneous Poisson process with a homogenous Gaussian rate function *ɷ*(*t*) centered at *μ_i_* with standard deviation *σ_i_* The time of maximum firing rate for the *i^th^* GC, *μ_i_*, is specified with respect to the time of *Ready.*

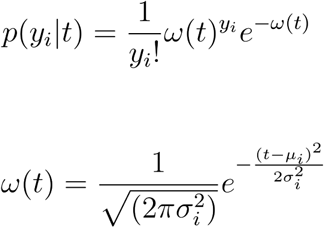

Due to scalar variability, the internal estimate of elapsed time 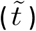 has a probabilistic relationship to the objective elapsed time (*t*). We formulated this relationship as a conditional Gaussian probability distribution whose mean is (*t*), and standard deviation scales with (*t*) by a scaling factor *ɷ_b_* This scaling factor is analogous to the Weber fraction introduced for the behavioral modeling. To model the basis set in the presence of this variability, we derived the expectation of the basis set across trials as a function of elapsed time, *t*, by marginalizing over this distribution.

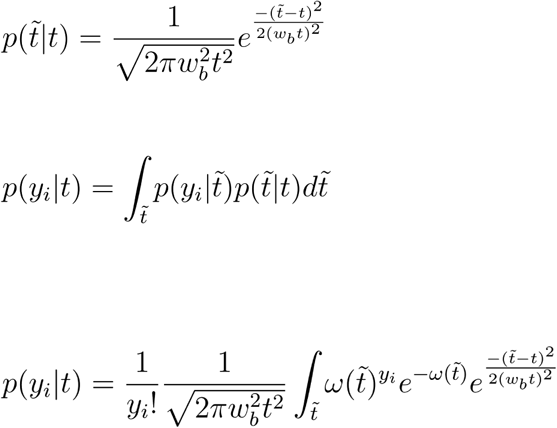

This transformation results in two forms of inhomogeneity across the basis set kernels: (1) it reduces the amplitude of kernels as a function of time, and (2) it causes kernels to become wider as a function of time (Supp. Fig 1). As expected, inferring elapsed time from this perturbed basis set using a maximum likelihood decoder produces scalar variability (Supp. Fig. 1).

We introduced a simplified parameterization to capture these two inhomogeneities. The reduction in amplitude was modeled by a decaying exponential function, *g*(*t*), with time constant *τ_basis_*, and the increase in width was modeled as a linear function, 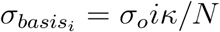, where *i* indexes neurons ordered according to their preferred time interval, *N* is the total number of neurons (*N* =500) and *k* is the proportion of increase in the width *σ_o_*. A detailed analysis of the robustness of model predictions upon varying these parameters can be found in Supp Fig. 3. The resulting function that describes the rate of the *i^th^* GC is:

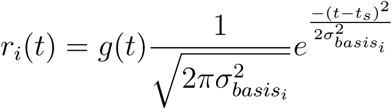

where

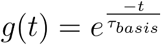

In the model, the PC activity is computed as a weighted sum of GC activity.

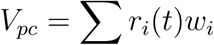

where *ɷ_i_* represents the synaptic weight of the *i^th^* GC.

Similar to previous work^32,120^, LTD in TRACE is modeled for each GC-PC synapse as proportional to the rate of firing of respective GCs shortly before the firing of climbing fibers (CFs) at the time of *set* The time before CF firing at which GC-PC synapses become eligible for LTD is called the eligibility trace (**ϵ**)^42,73^, which we assume occurs 50 ms before *Set* in the model. In the absence of CF stimulation and in the presence of GC firing, a weak restoring force (LTP) acts to reverse learning. The dynamics of LTD and LTP were governed by their respective time constants, ^-^*τ_ltd_* and *τ_ltp_*. A more detailed analysis of variation of these parameters can be found in Supp. Fig 4. In the absence of learning, synapses would gradually drift toward the baseline, *ɷ_o_.*

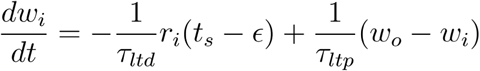

The presence of the eligibility trace implies that any CF firing at the onset of will have no bearing upon the plasticity of the GC-PC synapses. Similarly, GC activation at the time of *Set* will be irrelevant to learning of the prior. Further, our results remain qualitatively unchanged under assumptions of more complex functions of the eligibility trace.

Purkinje cells (PCs), which constitute the sole output of the cerebellar cortex are inhibitory. Since LTD reduces PC output, LTD-dependent learning has a net excitatory effect on the membrane potential of dentate neurons *V_dn_*. Furthermore, individual neurons in the dentate nucleus (DN) receive distal synaptic input from both extra-cerebellar afferents, mossy and climbing fibers. We assume this effective input to be constant (*I_eff_*). Finally, large columnar DN neurons are rich in axon collaterals that make autaptic connections^44^. With these elements, the membrane potential of DN neurons can be modeled as:

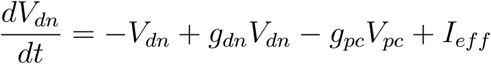

where *g_dn_* and *g_pc_* correspond to conductances associated with the autaptic input and PCs, respectively. For the simulations in the main text, we set *g_dn_* to 1, which corresponds to perfect integration, and set *I_eff_* to a fixed value equal to the average PC activity so that DN neurons receive similar levels of excitation and inhibition. However, TRACE exhibits robust Bayesian behavior under significant variation of both *I_eff_* and *g_dn_* (Supp Fig. 2). With these assumptions, the model can be simplified as follows:

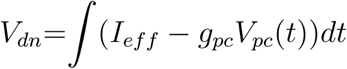

We simulated trial-by-trial dynamics by generating a spiking model for TRACE. On each trial and for each GC, we generated spike-trains according to a non-homogenous Poisson process whose rate was specified by the corresponding kernel in the basis set. Spikes were convolved with an excitatory postsynaptic gaussian kernel with standard deviation 20 ms. GC-PC synapses underwent LTD according to the level of activity of corresponding GCs at the time e (eligibility trace) before the time of *Set.*

#### Parameters values

The steady state behavior of TRACE is primarily governed by the parameters of the basis set and its dynamics by the learning rate parameters. The basis set has three parameters; *τ_basis_*, specifying attenuation, and *σ_o_*, *κ*, specifying increase in width. In Figure 1-6, the basis set parameters were *σ_o_* = 100 ms, *κ* = 0.2 and *τ_basis_* = 1000 ms. The TRACE model can tolerate a wide range of variation in these parameters (Supp. Fig. 3).

The learning dynamics in TRACE are characterized by changes in the magnitude of synaptic weights (*A_eff_*) and an effective time constant (*τ_eff_*) needed to reach such magnitudes. These variables are controlled by the *τ_ltd_* and *τ_ltp_* and the width of the prior distribution. In Supp. Fig. 4, we show how *A_eff_* and *τ_eff_* vary with *τ_ltd_*, and *τ_ltp_*. In Figure 5b, we used the prior distributions used in a previous study ^13^ and report the average deviation (across 100 simulations) of the PC activity in bins of 20 trials with *τ_ltd_* = 100 and *τ_ltp_* = 300 ms. Synaptic weights were not allowed to fall below zero and the lowest permissible value for long-term depression was *T_ltd_* = 50 ms.

In Figure 7a, which shows the TRACE behavior with all components intact, we use the same parameters as those used for Figures 1-6-c. In Figure 7b, we vary the basis set parameters. There is no exponential decay in Figure 7b and no widening of sigma. Original kernel width remains *σ_o_* = 100 ms. In Figure 7c, we set *τ_basis_* = 400 ms. All other parameters remain unchanged. In Figure 7d, we remove the learning equation from the model and do not make any adjustment to *V_dn_*. In Figure 7e, we remove the integration component of the model.

## Acknowledgements

The authors would like to thank Greg Horowitz, Seth Egger and Evan Remington for comments on an earlier version of the manuscript. D.N is supported by the Rubicon Grant (2015/446-14-008) from the Netherlands Scientific Organization (NWO). M.J. is supported by NIH (NINDS-NS078127), the McKnight Foundation, the Sloan Foundation, the Klingenstein Foundation, the Simons Foundation, the Center for Sensorimotor Neural Engineering, and the McGovern Institute

---

**Supplementary Figure 1:**
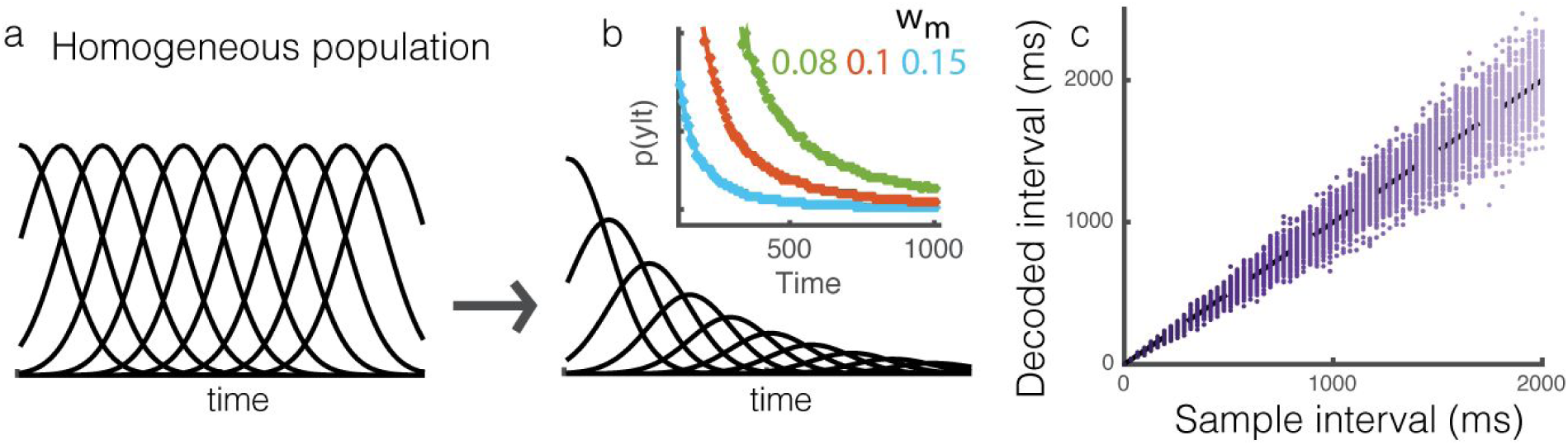
Probabilistic motivation of the basis set a) A homogeneous basis set in the absence of scalar noise (sub-sampled), b) The expected profile of the same basis set under the influence of scalar noise. This causes the expected basis set to attenuate and become wider with time. The inset shows the profile of attenuation for three weber fractions (w_m_). c) A maximum likelihood estimate of time decoded from the activity of the modified basis set expresses scalar variability.

**Supplementary figure 2.**
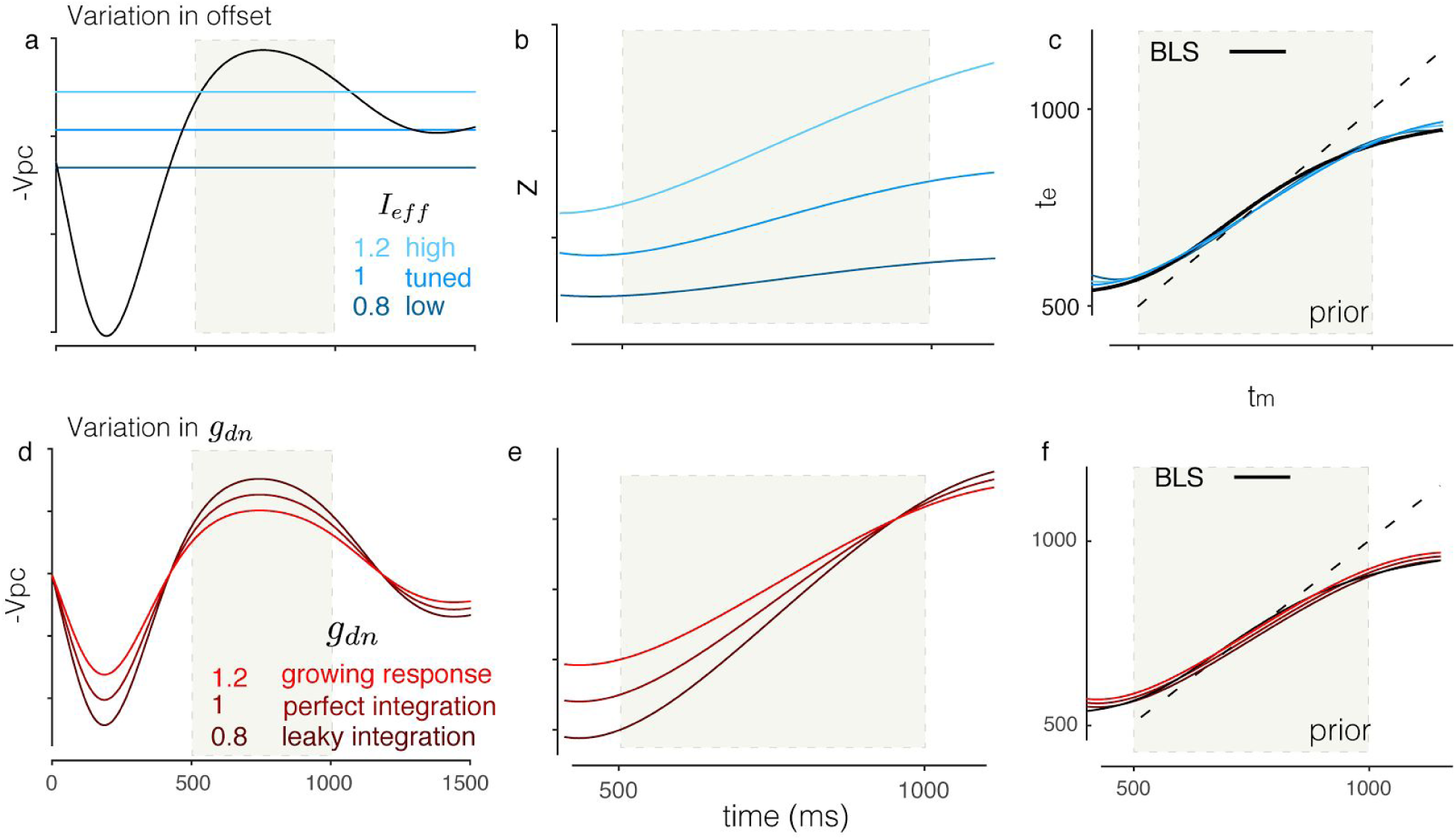
Variation of the parameters of integration in TRACE, a-c) The effect of varying I_eff_ (shades of blue) on model behavior, a) −V_pc_ as a function of time (black) with the three offset levels overlaid, b) The integrated output with the three offsets, c) Linearly transformed output for the three offsets. The black shows the Bayes least-squares function; i.e., Bayesian interval estimate (t_e_) as a function of the measured interval (t_m_). The dark to light blue correspond to cases when the offset is 20% lower (low), equal (tuned), or 20% higher (high) than the mean of −V_pc_. d-f) The effect of varying g_dn_ (shades of red) on model behavior. Panels follow the same organization as the top row. The dark to light red correspond to a leaky integration (g_dn_ = 0.8), perfect integration (g_dn_ = 1), and a case where responses grow faster than perfect integration (g_dn_ = 1.2). If g_dn_ is significantly greater than 1, the neuron becomes unstable and the model breaks down. The shaded area in all panels correspond to the support of the prior distribution.

**Supplementary figure 3.**
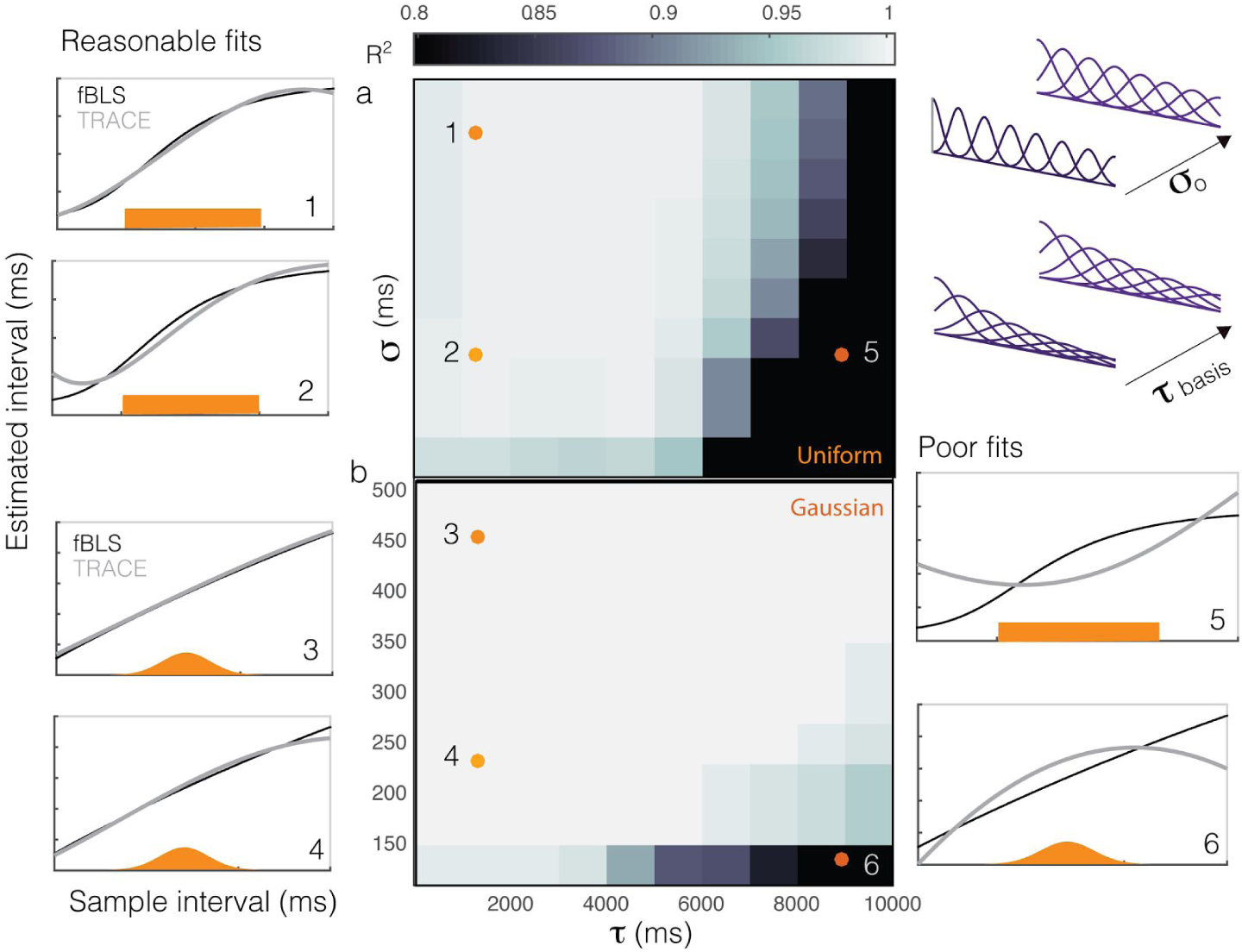
Robustness to model parameters. A grayscale heotmop showing the goodness of fit between the TRACE model (gray line) and Bayes least-squares (BLS - block line) while varying parameters of the basis set (kernel width σ_o_ and time constant of exponential decay τ_basis_) for a) uniform and b) Gaussian prior distributions (oronge). Plots on the left (1-4) and right (5-6) show examples with high and low R^2^, respectively.

**Supplementary figure 4.**
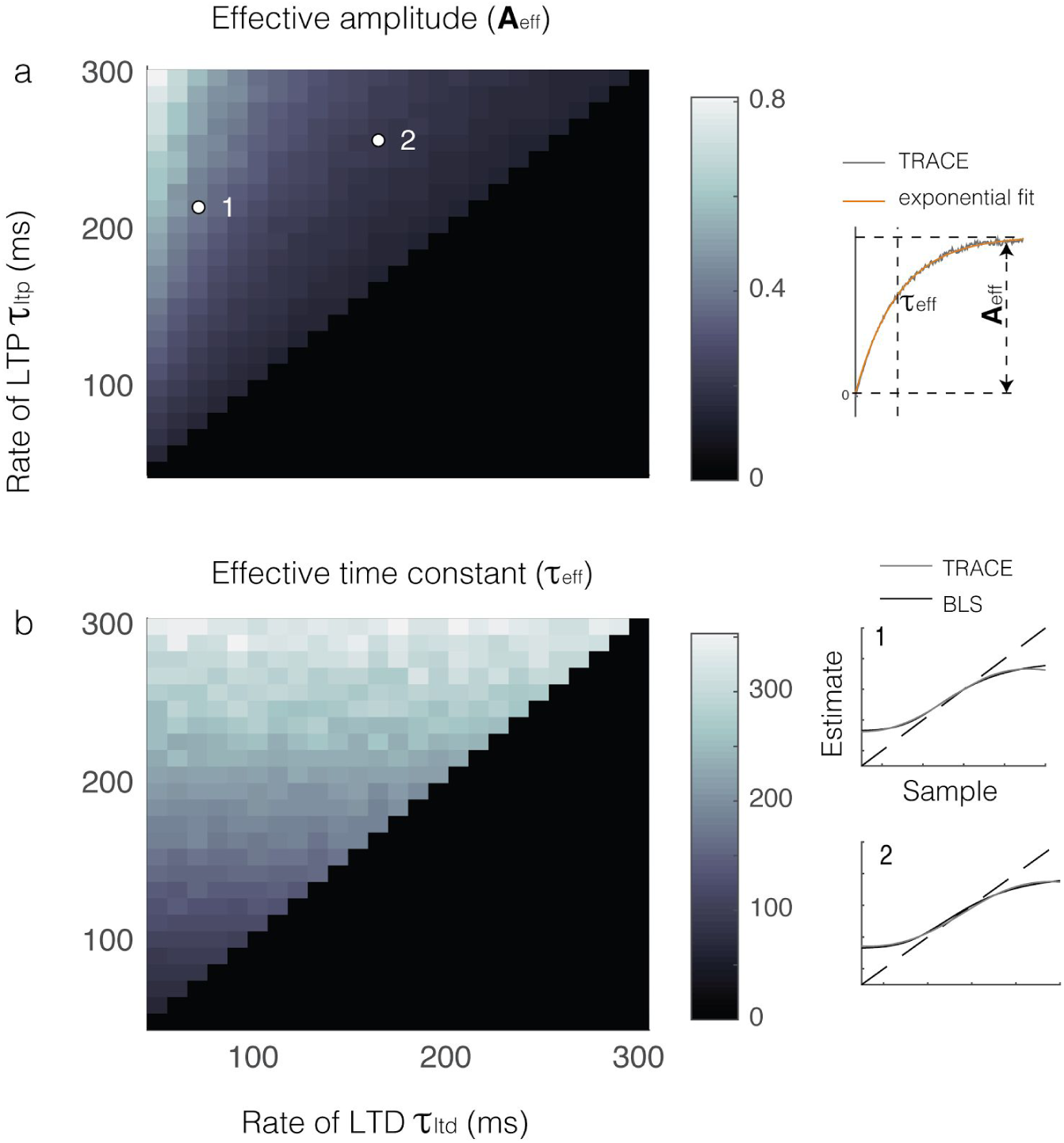
Learning dynamics, a) Grayscale indicates the effective amplitude (A_eff_) of synaptic weights across all neurons as a function of the time constants of LTD and LTP (τ_ltd_ and τ_ltp_). The inset shows how A_eff_ and τ_eff_ were determined from sum of synaptic weights across neurons over 200 observations. Inset below shows linearly transformed integrated TRACE outputs (gray) compared to α Boyes least-squares function (block) for two arbitrary points marked in a), b) Grayscale indicates effective time constant τ_eff_ as a function of τ_ltd_ and τ_ltp_.

**Supplementary Figure 5.**
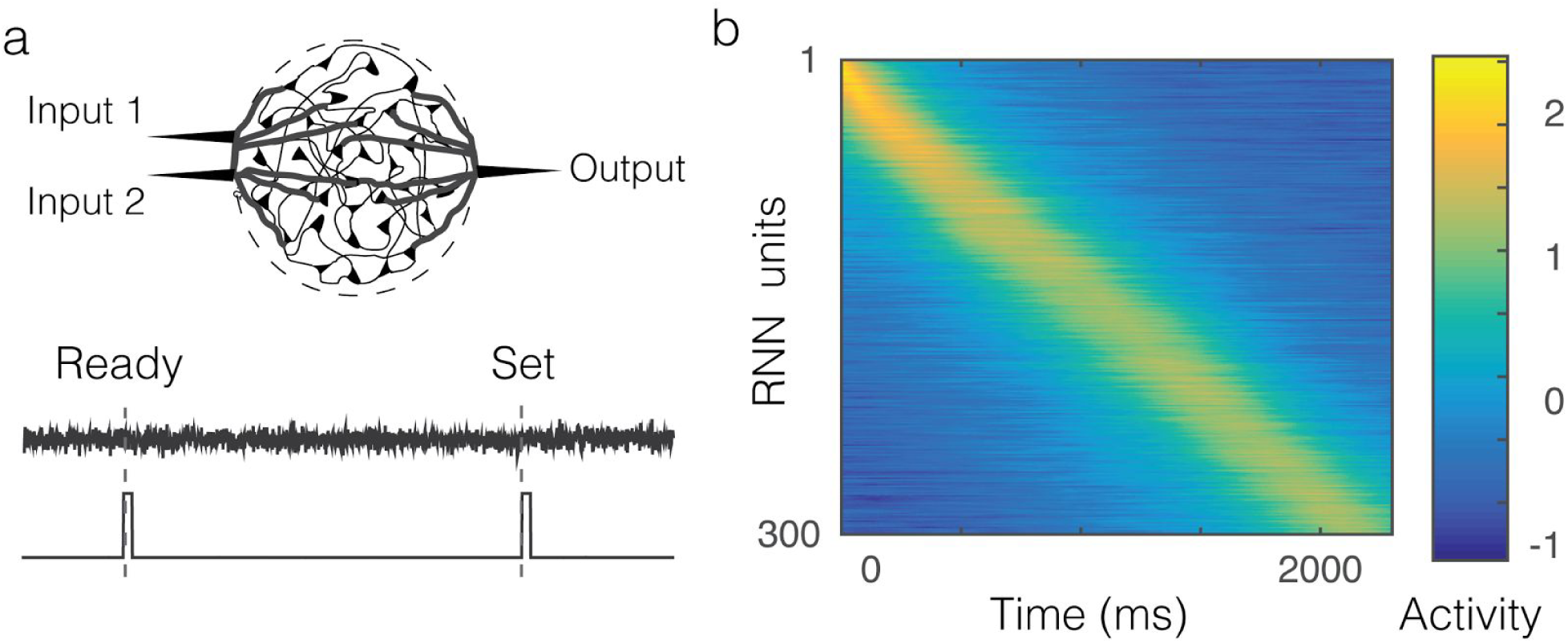
Demonstration of the effect of scalar variability on the basis set in a random recurrent network, a) a recurrent neural network driven by white noise and transient pulses associated with Ready and Set b) trial-averaged activity of all the units rank-ordered by peak activity as a function of time.

